# Virome profiling of *Culex tarsalis* through small RNA-seq: A challenge of suboptimal samples

**DOI:** 10.1101/2025.05.23.655811

**Authors:** Jaime Manzano-Alvarez, Sultan Asad, Duverney Chaverra-Rodriguez, Eunho Suh, Jason L. Rasgon

**Affiliations:** Department of Entomology, The Pennsylvania State University, University Park, PA, 16802, USA; Center for Infectious Disease Dynamics, The Pennsylvania State University, University Park, PA, 16802, USA; Dirección Académica, Universidad Nacional de Colombia Sede de La Paz, La Paz, Colombia; Department of Biochemistry and Molecular Biology, The Pennsylvania State University, University Park, PA, 16802, USA; The Huck Institutes of the Life Sciences, The Pennsylvania State University, University Park, PA, 16802, USA; The ONE Health Microbiome Center, The Pennsylvania State University, University Park, PA, 16802, USA; Institute of Energy and the Environment, The Pennsylvania State University, University Park, PA, 16802, USA

**Keywords:** Virome, RNA-Seq, sRNA, siRNA, piRNA, mosquito

## Abstract

Viral infections in mosquitoes trigger the RNA interference (RNAi) pathway, a key antiviral defense mechanism that generates virus-derived small RNAs (vsRNAs). Given the natural enrichment of vsRNAs during infection and their stability, small RNA sequencing (sRNA-seq) has emerged as a powerful tool for virome characterization. *Culex tarsalis* is a widely distributed mosquito species in North America and is an important vector of West Nile virus (WNV). Previous studies have shown that co-infection with insect-specific viruses (ISVs) can modulate WNV replication in *Cx. tarsalis*, highlighting the importance of characterizing the virome of this species. Here, we investigated the virome of *Cx. tarsalis* populations across 5 states of the Midwestern United States using sRNA-seq. We analyzed samples from 17 geographic locations which were collected under suboptimal field conditions during the COVID-19 pandemic, presenting challenges related to sample integrity. Despite these challenges, sRNA-seq proved to be a reliable method for virome analysis. We identified seven viruses associated with *Cx. tarsalis*, along with their respective sRNA (siRNA and piRNA) profiles. These findings not only deepen our understanding of ISVs, but also demonstrate the utility of sRNA-seq in non-ideal situations, enabling the collection and analysis of samples under real-world surveillance scenarios.

## Introduction

Insect specific viruses (ISVs) are viruses that infect insects, but which do not infect vertebrate animals. Due to host restriction, they have gained attention as potential biocontrol agents and as tools for studying vector biology, as well as potential agents for disease control [1–4]. In the case of mosquitoes, their virome includes ISVs and pathogenic arboviruses, the latter being able to infect vertebrates as well. Mosquito ISVs can interact with pathogenic arboviruses such as West-Nile virus (WNV), the most important arbovirus in the United States [5,6]. Eilat virus, an ISV naturally found in *Culex univittatus, Cx. pipiens* and *Anopheles coustani* [7–9], can induce superinfection exclusion against WNV in *Cx. tarsalis* mosquitoes [3], highlighting the public health significance of studying ISVs in this mosquito species. More than 39 ISVs have been discovered in *Cx. tarsalis* samples, but most virome studies for this species are restricted to California, [10]. Consequently, even though this species is distributed across North America, its virome remains unexplored across the majority of its range.

sRNAs are short, non-coding RNA molecules that can play critical roles in gene regulation and antiviral defense mechanisms [11]. In mosquitoes, sRNAs are key players in the RNA interference (RNAi) pathway, an important component of the innate immune system that targets and degrades viral RNA. Some classes of sRNAs that participate in RNAi viral response, include small interfering RNAs (siRNAs; 21nt), and piwi-interacting RNAs (piRNAs; 24-30nt), which are involved in different biogenesis pathways and have different biological functions [12–15]. The sRNAs produced during RNAi response against viral infection are considered virus-derived small RNAs, and are the focus of virome studies because they provide a molecular signature of viral presence. Thus, the application of sRNA-seq in mosquito virome studies allows for the identification of known and novel viruses [16–19], as well as a unique virus molecular signatures, or sRNA profiles, which could be used for identification and understanding of their interactions within their host (reviewed in [20]). Importantly, since small RNAs (sRNAs) are naturally enriched during viral infections in insect cells, the use of sRNA-seq has proven to be an efficient, specific and reliable approach in previous studies[16– 18,20–22]. While both RNA-seq and sRNA-seq have been used for virome identification in mosquitoes [2,10,16,20,21], sRNA-seq has advantages due to the natural enrichment of sRNAs during viral infections, as well as their higher stability due to their small size, association with proteins, and 2′-O-methylation at the 3′ terminal which protects them from degradation by exonucleases [20,23–25]. As a result, sRNAs in a sample may maintain higher integrity over time compared to longer RNAs, such as mRNA.

The Covid 19 pandemic mitigation strategies caused a wide range of challenges for researchers, impacting both field and lab work [26], tremendously limiting our ability to collect samples in ideal conditions of RNA integrity preservation. To address this challenge, we collaborated with mosquito control agencies that routinely collect mosquitoes for surveillance purposes across multiple locations in the United States. Since trapping and shipping methods do not always allow for strict adherence to cold-chain protocols, preserving RNA integrity can be difficult. Given that sRNAs are more stable than total RNA and are naturally enriched during viral infections, we used a sRNA-seq approach. In this study, we investigated the virome of *Cx. Tarsalis* mosquitoes collected from 17 geographic locations across five Midwestern states using sRNA sequencing and bioinformatic analyses.

## Methods

### Sample Collection

Wild *Cx. tarsalis* mosquitoes were collected from 17 different locations across 5 states in the USA (CA, CO, TX, WA, UT; 3 locations per state except for CA and TX with 5 and 2 locations respectively). Mosquitoes were trapped using CDC and BG traps, then identified and immediately stored in DNA/RNA shield (Zymogen Cat. No. R1200) by Vector Disease Control International (VDCI), and local mosquito control agents. Mosquitoes were shipped at room temperature to the Rasgon Lab at The Pennsylvania State University, where they were sexed and immediately frozen at -80°C until further processing. Additionally, virus-free *Cx. tarsalis* mosquitoes (KNWR strain) from the Rasgon lab were processed as a control.

### RNA Extraction and small RNA sequencing

RNA was extracted from pools of 3 female *Cx. tarsalis* mosquitoes per geographic location using Qiazol lysis Reagent (Qiagen Cat. No. / ID: 79306) and Direct-zolTM RNA MicroPrep kit (Zymogen Cat. No. R2062). Briefly mosquitoes were thawed in DNA/RNA shield solution and were rinsed once in 1xPBS. 1000ul of Qiazol and 0.2-0.5 nm metallic beads were added to the pools in 1.5ml tubes, then samples were homogenized using Qiagen tissue lyser for 90 second at 30.0 hertz/second, and then centrifuged at 15000 xg for 1 minute. The clear lysate was transferred to a new 2ml tube and RNA was purified using Direct-zolTM RNA MicroPrep kit using manufacturer instructions. Extracted RNA was sent to Novagene USA to perform small RNA sequencing using NovaSeq SE50 sequencing.

### Small RNA reads preparation

Raw single end sequences were trimmed using cutadapt (v5.0), removing reads with average Phred quality below 20, ambiguous nucleotides, and reads with length shorter than 15nt and/or without adaptors. The remaining reads were mapped to the *Cx. tarsalis* reference genome CtarK1 [27] using bowtie2 (version 2.5.1), and bacteria reference genomes via Kraken2 (version 2.1.3). Unmapped reads were then considered as processed reads and were used for further steps.

### Contig assembly and extension

Processed reads that were 20-30nt long were used for contig assembly in each library (pool per location) using Velvet (version 1.2.10) and Spades (version 3.13.0), as previously described [21]. Contigs >200nt from each location were grouped using CD-HIT (version 4.8.1) requiring 90% of coverage with 90% of identity in order to remove redundancy and compared against the NCBI NT and NR databases with BLAST. Contigs that were identified as viral or unknown origin were chosen, and then sRNA profiles and coverage of each contig were plotted using in-house Perl v5.16.3, BioPerl library v1.6.924 and R v4.3.2 scripts, as previously described [16]. Contigs were manually curated by selecting those that showed a 21nt peak in their sRNA profiles, indicative of Dicer-processed sRNA species. Contigs with mapped reads displaying even coverage in both forward and reverse orientations were selected. Filtered contigs from every location were properly labeled and grouped together with CD-HIT to designate representative contigs. Contigs were then mapped against the processed reads for co-occurrence analyses based on hierarchical clustering with Pearson correlation using R scripts. Contigs that clustered together were used for contig extension in Spades as trusted contigs with all the libraries that mapped against them (minimum of 300 reads). All the extended contigs were grouped together and then manually curated. Contigs that had their top three BLAST matches against viral sequences of the same species were used to identify their likely origin. Finally, reference genomes of the identified viruses were subsequently used as templates for sRNA profiling, genome coverage and co-occurrence analyses as previously described.

### Data reporting

RNA libraries are deposited at https://doi.org/10.5281/zenodo.15412939

### Use of AI

ChatGPT and GitHub Copilot were used to assist in generation of some of the pipeline code used for data analysis. Generated code was tested, debugged, and validated manually. All codes and scripts are available at https://github.com/Zureishon/Virome.

### Figure generation

Graphs and plots were made with R Studio (2023.3.0.386; PBC, Boston, MA, USA) and Biorender.com. Final figures were assembled using Adobe Illustrator 2023 (27.4.1; Adobe, San Jose, CA, USA). Statistical summaries of RNA libraries and assembled contigs are provided as supplemental material.

## Results

### Seven ISVs Identified in Culex tarsalis via sRNA-Seq

Pools of three unfed female *Cx. tarsalis* mosquitoes from each of the 17 locations were processed for RNA extraction and sRNA-seq. Due to suboptimal collection conditions for RNA preservation, challenges in RNA extraction and sequencing were anticipated. Post-trimming, library sizes varied between 3–25 million (M) reads, reflecting differences in sRNA availability across samples. Most libraries contained fewer than 2M reads within the target sRNA size range (20–29 nt), with the exception of UT1, UT2, and UT3, which exceeded 4M target reads (SFig. 1a). Reads were first mapped to the *Cx. tarsalis* genome and bacterial sequences for filtering. In most libraries, over 80% of reads mapped to the *Cx. tarsalis* genome; however, in CO2, WA1, WA3, and TX2, less than 45% mapped, suggesting higher proportions of non-host sequences (SFig. 1b). Given the limited size of the libraries, we employed two assembly strategies using Velvet and SPAdes to maximize the contig assembly. We identified 99 contigs longer than 200 bp, which were subsequently BLASTed to determine their origins. Contigs of viral origin were detected in 5 out of the 17 sampled populations (Fig. 2a). To refine our virome analysis, we examined sRNA profiles and coverage maps of contigs classified as viral or of unknown origin. Contigs displaying a 21-nt peak and even coverage were used for further analysis. From this selection, 26 representative contigs were chosen for co-occurrence analyses based on their RPKM values, which grouped samples into 8 distinct clusters (SFig. 2). These clusters, along with their associated representative contigs, were then subjected to a second round of contig assembly using SPAdes. This approach yielded 131 unique contigs longer than 200bp with a maximum of 1387bp, of which 65 contigs were identified as viral in origin, corresponding to seven ISVs. To determine the closest viral references, all contigs were BLASTed, and available reference genomes were used for co-occurrence analyses (Fig2b). The most widely distributed viruses were Marma virus (MV) and *Culex* narnavirus 1 (CN1), present across all five states. *Culex* Bunyavirus 2 (CB2) and Wuhan Mosquito virus 6 (WMV6) were detected in four states, while Partitivirus-like *Culex* mosquito virus (PCMV) was found in three states. Lastly, *Culex* Iflavi-like virus 4 (CIV4) and Hubei mosquito virus 4 (HMV4) were identified in two states (Fig. 2b). All detected viruses were classified as RNA viruses (Table 1), all of which have been previously reported in *Cx. tarsalis* mosquitoes from other areas [28]. No viral reads were detected in the KNWR insectary sample thus it was excluded from subsequent analyses.

**Table 1.**
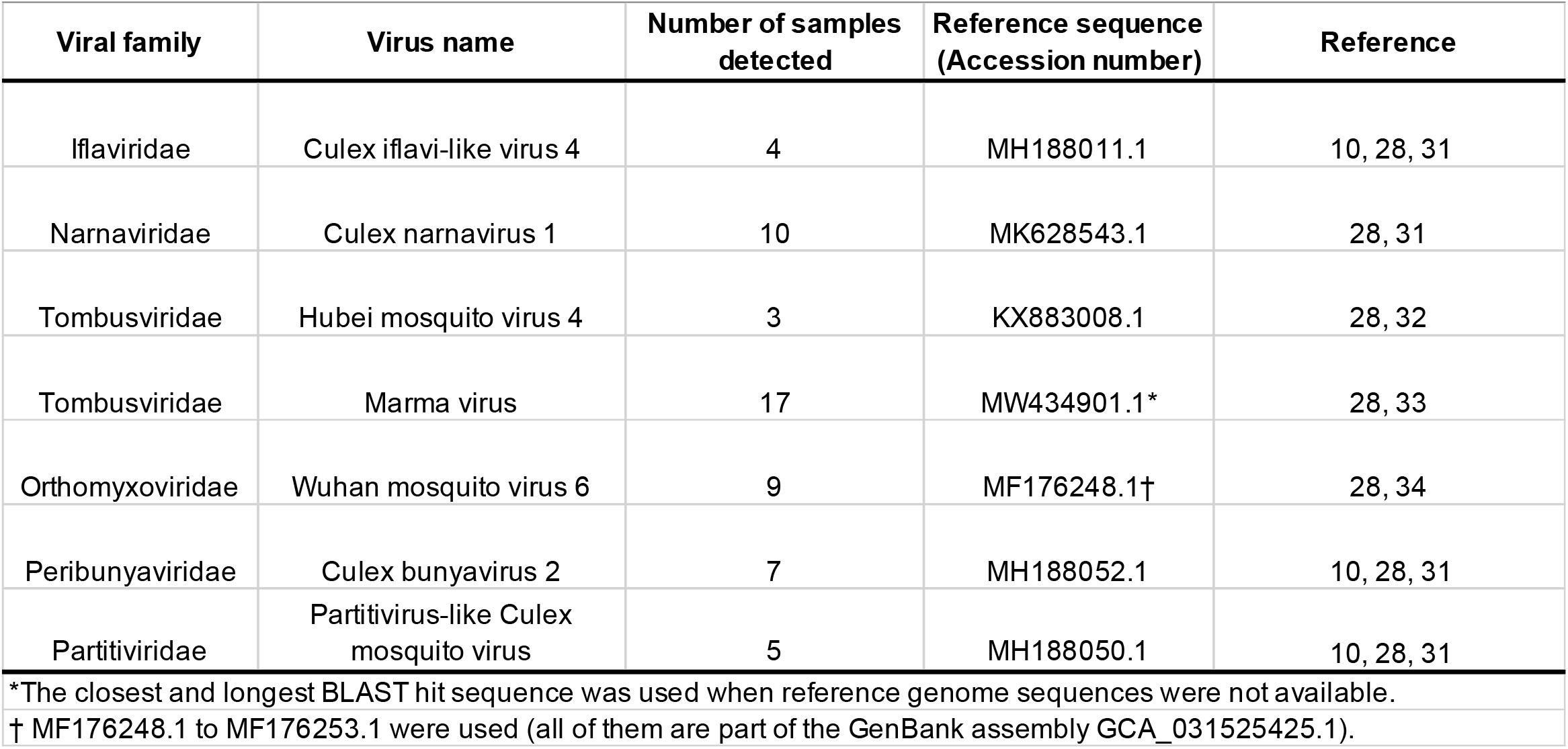
Classification and previous reports of characterized viruses in North America. GenBank accession numbers refer to the viral reference sequences used for the assembly of sRNA profiles.

**Figure 1.**
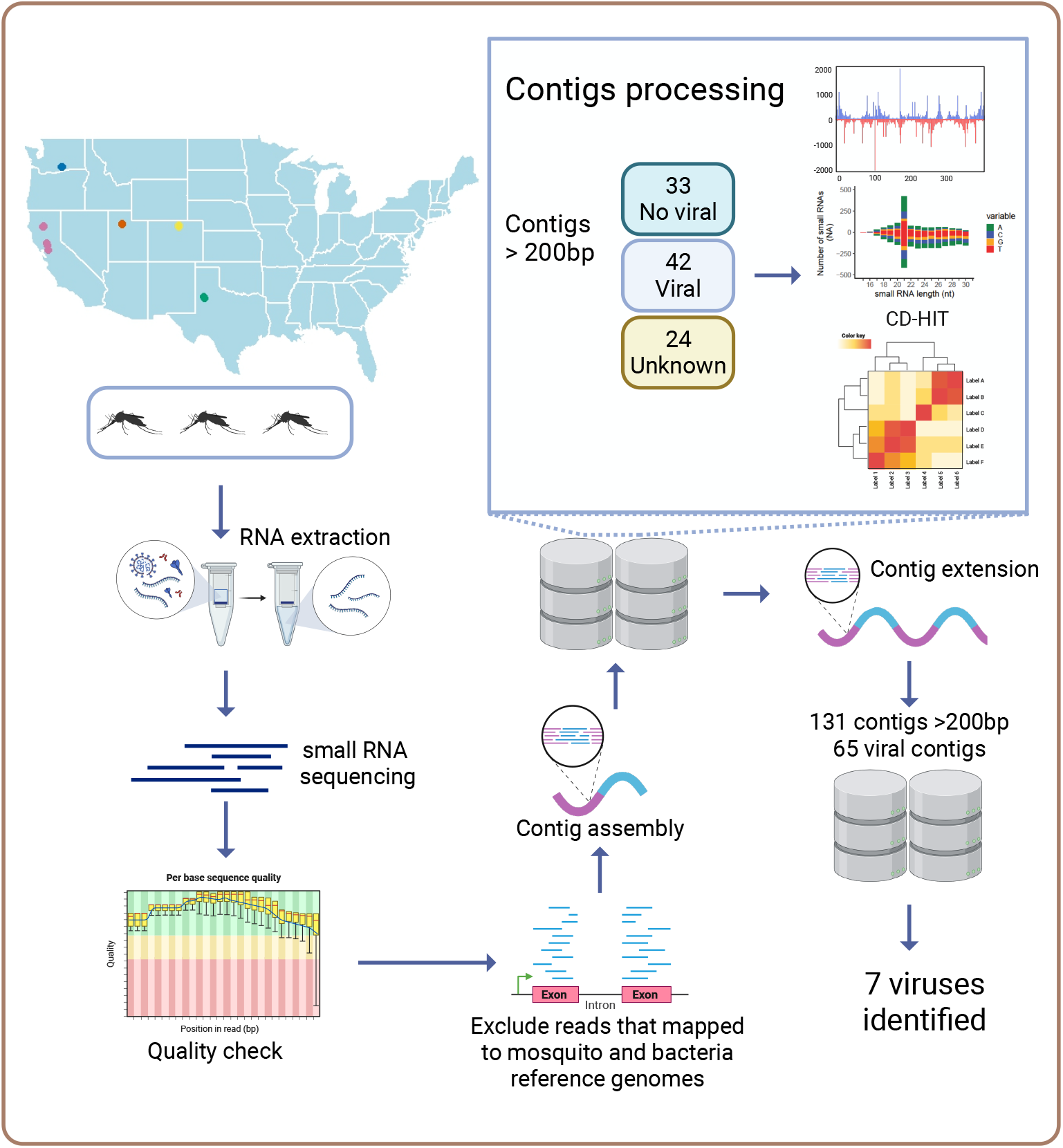
Graphical methods of metagenomics pipeline for virome profiling. Pools of 3 female mosquitoes per location were used for RNA extraction and sRNA-seq. Quality check steps were done in raw reads, then used for contig assembly and processing. Viral and unknown contigs were used for co-occurrence analyses and contig extension. Extended contigs were BLASTed for virus identification, and available viral reference sequences were used for virus co-occurrence analyses and sRNA profile design.

**Figure 2.**
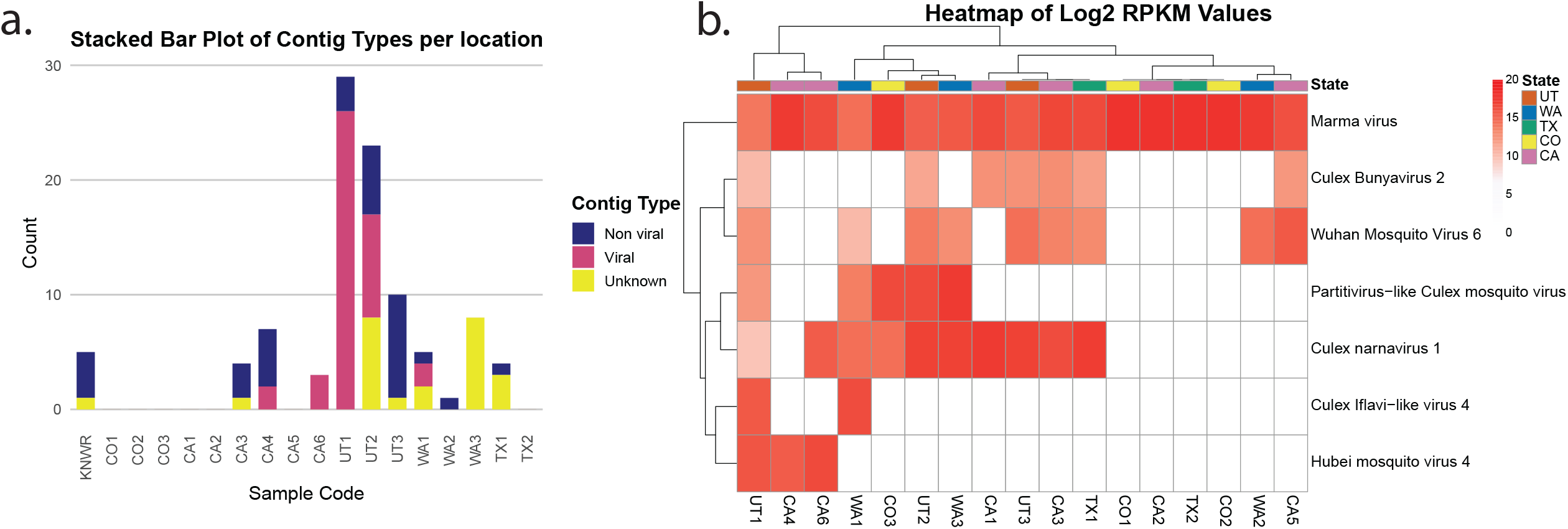
(a) Contig classification per sample and ISV distribution. For each library, assembled contigs longer than 200 bp were BLASTed and classified according to their origin. (b) Heatmap shows the Log_2_ RPKM values for each library, based on mapping them to the reference genomes of the ISVs identified through metagenomics analyses.

We then analyzed sRNA profiles and coverage maps for each identified virus. sRNA profiles are histograms displaying the absolute number of reads per size that align to the genome in the forward and reverse orientations; piRNA profiles follow the same principle but only include reads enriched for U at the 5’ end or A at the 10th nucleotide. Coverage maps illustrate the depth of alignment by showing how many times each nucleotide in the genome is covered by a read, plotted separately for 21-nt reads and 24–29-nt reads. Most of the identified viruses showed a peak in the 21-nt size reads, indicative of siRNA pathway activation, with participation of the piRNA pathway represented by the presence of 24-30nt reads. However, CIL4 and HMV4 exhibited an enrichment of sRNA reads mostly in the forward sense, with almost identical sRNA and piRNA profiles. Coverage maps revealed that sRNAs were mapped across most of each viral genome, except for HMV, which displayed regions with little to no read coverage (Fig. 3).

**Figure 3.**
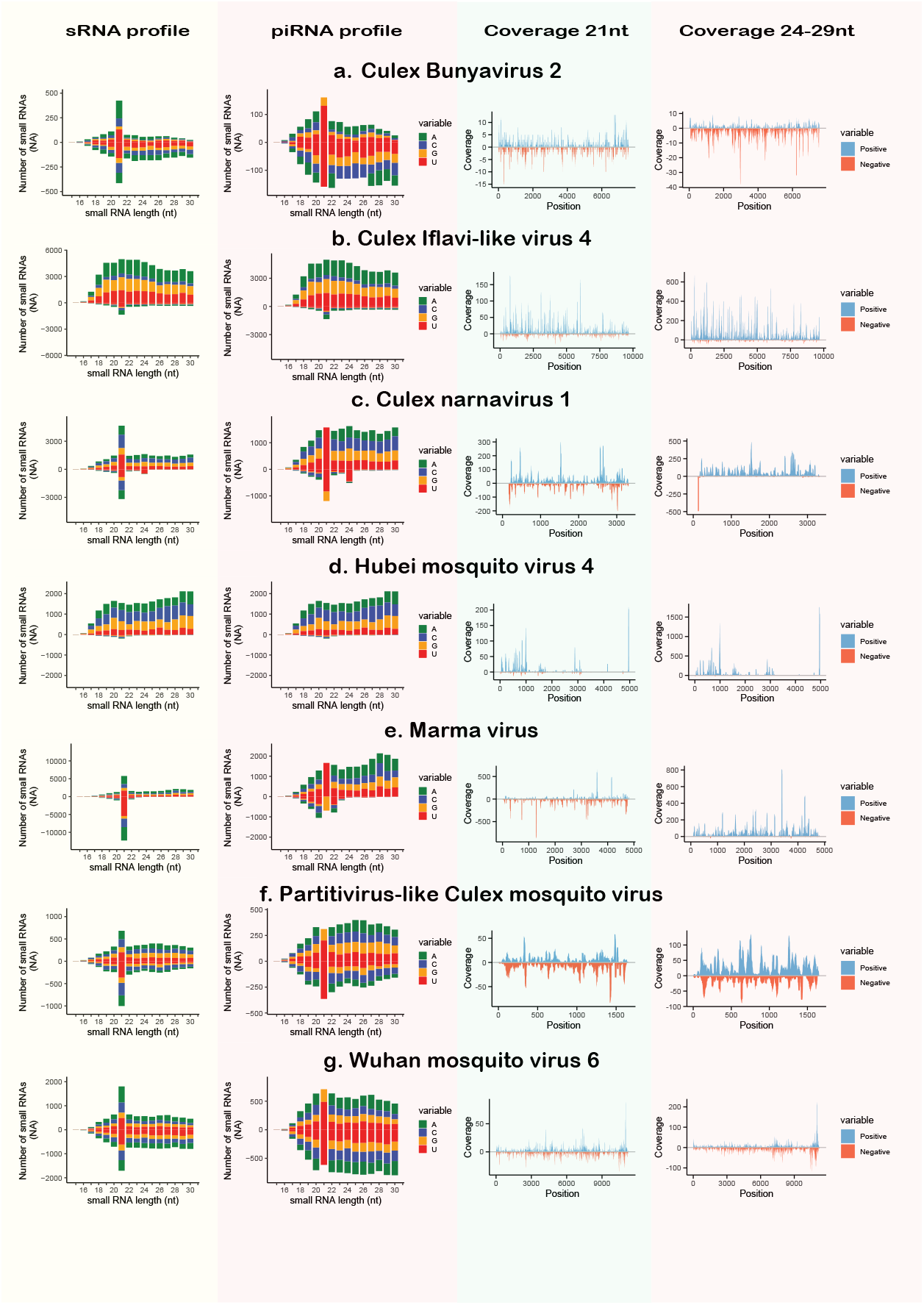
sRNA profiles and coverage plots of the ISVs. All libraries were used to generate these graphs. sRNA profiles are histograms representing the frequency of mapped read sizes, with colors indicating 5’ nucleotide bias. piRNA profiles are sRNA profiles filtered to include only reads with a 5’ U bias or an A at the 10th nucleotide. Coverage plots were generated using reads of either 21 nt in length, representing siRNAs, or reads of 24–29 nt, representing piRNAs. Colors in the coverage plots indicate whether the reads aligned to the sense (blue) or antisense (red) strand of the reference viral genomes.

## Discussion

RNA-seq has become a powerful tool for the surveillance and discovery of new viruses in insects. In mosquitoes, its use has been demonstrated across several countries, especially in species of the genus *Aedes* [16,18,20–22,29,30]. In North America, some studies have analyzed the virome of *Culex* mosquitoes using either long or small RNA-seq, with most focusing on Canada, and the state of California in the USA. The majority of these studies have examined *Cx. pipiens* or *Cx. quinquefasciatus*, with only two investigations including *Cx. tarsalis* [10,28,31,32]. In this study, we detected 7 of the 39 viruses associated with *Cx. tarsalis* from California [10].

The use of sRNA-seq enables the exploration of the sRNA profiles, which can reflect how viruses interact with the mosquito immune response. In most cases, the sRNA profiles in this study resemble previously described profiles for each virus [31], except for HMV4, and PCMV. These results support the reliability of sRNA profiles, even under potentially different environmental and physiological conditions. Importantly, we observed the activation of the siRNA pathway in response to most viruses. This is expected since the siRNA pathway is typically triggered by viral double-stranded RNA, cleaving it into 21-nt siRNAs that guide the degradation of viral genomes [33,34]. In contrast, HMV4 and CIV4 elicited a stronger piRNA pathway response compared to siRNA. The piRNA pathway primarily originates from the mosquito’s own genome, producing slightly longer sRNAs (24–30 nt) from piRNA clusters [12,35,36]. In most cases, these clusters are transcribed from a single DNA strand, leading to a characteristic strand bias in piRNAs [37,38]. Activation of those pathways have been inferred for several viruses infecting *Culex* mosquitoes, both *in vivo* and *in vitro* [39], as well as in virome-wide analyses via sRNA-seq [31]. Coverage plots indicate that most sRNAs mapping to HMV4 and CIV4 were strand-biased, aligning predominantly to a single strand, consistent with a piRNA response with impairment of siRNA response (e.g. Dicer inhibition) [20,33].

An important aspect of this study is that mosquitoes were collected by mosquito control agencies as part of their routine surveillance, using overnight traps, during an active pandemic that limited ability to ship or process samples in a timely manner. Under these conditions, it is difficult to guarantee high-quality RNA, as mosquitoes may die before collection or processing is complete, leading to RNA degradation. To overcome this, we took advantage of the higher stability of small sRNAs. The sequencing sRNA libraries generated in this study were, in most cases, around five times smaller than those used in previous work [16,22,30]. As a result, the assembled viral contigs were relatively short, limiting our ability to reconstruct full viral genomes. Nonetheless, we were able to accurately identify viral infections, highlighting the potential of sRNA-based approaches for RNA virus surveillance in mosquitoes (or other insects) even when collection conditions are suboptimal. Improvements for further studies could include better coordination with mosquito control agencies to enhance sample preservation, as well as the adoption of more efficient RNA extraction methods. These adjustments could significantly improve RNA quality and increase the likelihood of assembling complete viral genomes, capture more viral diversity, or the identification of novel viruses.

In conclusion, this study demonstrates the utility of sRNA-seq for virus surveillance in *Cx. tarsalis*, providing new insights into the mosquito anti-viral immune response. Despite challenges related to sample integrity, we observed activation of both siRNA and piRNA pathways in response to most viruses, with HMV and CIV4 triggering only a piRNA response. Moving forward, expanding sample collection efforts and refining preservation and extraction methods will strengthen future virome surveillance studies.

## Acknowledgements

We thank all the mosquito control agencies and researchers who collected and provided an initial mosquito species identification: Tooele Valley Mosquito Abatement District, Benton County Mosquito Control District, Dr. Corey Brelsfoard from Texas Tech University, VDCI Mosquito Management, The Mosquito and Vector Control Association of California (MVCAC), and Sacramento-Yolo Mosquito and Vector Control District.

J.M.A. was supported by the Fulbright Pasaporte a la Ciencia program, a collaboration between Fulbright Colombia and ICETEX, as part of the Colombia Científica program in response to the “foco-reto país en Salud”. Funding for this work was provided by NIH grant R01AI150251, USDA Hatch Project 4769, and funds from the Dorothy Foehr Huck and J. Lloyd Huck endowment to JLR.

## References

1. Agboli E, Leggewie M, Altinli M, Schnettler E. Mosquito-Specific Viruses— Transmission and Interaction. Viruses 2019, Vol 11, Page 873. 2019;11: 873. doi:10.3390/V11090873

2. de Almeida JP, Aguiar ER, Armache JN, Olmo RP, Marques JT. The virome of vector mosquitoes. Curr Opin Virol. 2021;49: 7–12. doi:10.1016/J.COVIRO.2021.04.002

3. Joseph RE, Bozic J, Werling KL, Krizek RS, Urakova N, Rasgon JL. Eilat virus (EILV) causes superinfection exclusion against West Nile virus (WNV) in a strain-specific manner in Culex tarsalis mosquitoes. Journal of General Virology. 2024;105: 2017. doi:10.1099/jgv.0.002017

4. Romo H, Kenney JL, Blitvich BJ, Brault AC. Restriction of Zika virus infection and transmission in Aedes aegypti mediated by an insect-specific flavivirus. Emerg Microbes Infect. 2018;7. doi:10.1038/S41426-018-0180-4

5. Bolling BG, Olea-Popelka FJ, Eisen L, Moore CG, Blair CD. Transmission dynamics of an insect-specific flavivirus in a naturally infected Culex pipiens laboratory colony and effects of co-infection on vector competence for West Nile virus. Virology. 2012;427: 90–97. doi:10.1016/J.VIROL.2012.02.016

6. Goenaga S, Kenney JL, Duggal NK, Delorey M, Ebel GD, Zhang B, et al. Potential for Co-Infection of a Mosquito-Specific Flavivirus, Nhumirim Virus, to Block West Nile Virus Transmission in Mosquitoes. Viruses. 2015;7: 5801–5812. doi:10.3390/V7112911

7. Bennouna A, Gil P, El Rhaffouli H, Exbrayat A, Loire E, Balenghien T, et al. Identification of Eilat virus and prevalence of infection among Culex pipiens L. populations, Morocco, 2016. Virology. 2019;530: 85–88. doi:10.1016/J.VIROL.2019.02.007

8. Guggemos HD, Kopp A, Voigt K, Fendt M, Graff SL, Mfune JKE, et al. Eilat virus isolated from Culex univittatus mosquitoes from the Namibian Zambezi Region influences in vitro superinfection with alpha- and flaviviruses in a virus-species-dependent manner. PLoS One. 2024;19: e0312182. doi:10.1371/JOURNAL.PONE.0312182

9. Nasar F, Palacios G, Gorchakov R V., Guzman H, Travassos Da Rosa AP, Savji N, et al. Eilat virus, a unique alphavirus with host range restricted to insects by RNA replication. Proc Natl Acad Sci U S A. 2012;109: 14622–14627. doi:10.1073/PNAS.1204787109

10. Sadeghi M, Altan E, Deng X, Barker CM, Fang Y, Coffey LL, et al. Virome of >□12 thousand Culex mosquitoes from throughout California. Virology. 2018;523: 74–88. doi:10.1016/j.virol.2018.07.029

11. Gou LT, Zhu Q, Liu MF. Small RNAs: An expanding world with therapeutic promises. Fundamental Research. 2023;3: 676–682. doi:10.1016/J.FMRE.2023.03.003

12. Morazzani EM, Wiley MR, Murreddu MG, Adelman ZN, Myles KM. Production of Virus-Derived Ping-Pong-Dependent piRNA-like Small RNAs in the Mosquito Soma. PLoS Pathog. 2012;8: e1002470. doi:10.1371/JOURNAL.PPAT.1002470

13. Muellera S, Gausson V, Vodovar N, Deddouchea S, Troxler L, Perot J, et al. RNAi-mediated immunity provides strong protection against the negative-strand RNA vesicular stomatitis virus in drosophila. Proc Natl Acad Sci U S A. 2010;107: 19390–19395. doi:10.1073/PNAS.1014378107

14. Vodovar N, Bronkhorst AW, van Cleef KWR, Miesen P, Blanc H, van Rij RP, et al. Arbovirus-Derived piRNAs Exhibit a Ping-Pong Signature in Mosquito Cells. PLoS One. 2012;7: e30861. doi:10.1371/JOURNAL.PONE.0030861

15. Wu Q, Luo Y, Lu R, Lau N, Lai EC, Li WX, et al. Virus discovery by deep sequencing and assembly of virus-derived small silencing RNAs. Proc Natl Acad Sci U S A. 2010;107: 1606–1611. doi:10.1073/PNAS.0911353107

16. Aguiar ERGR, Olmo RP, Paro S, Ferreira FV, De Faria IJDS, Todjro YMH, et al. Sequence-independent characterization of viruses based on the pattern of viral small RNAs produced by the host. Nucleic Acids Res. 2015;43: 6191–6206. doi:10.1093/NAR/GKV587

17. Kreuze JF, Perez A, Untiveros M, Quispe D, Fuentes S, Barker I, et al. Complete viral genome sequence and discovery of novel viruses by deep sequencing of small RNAs: A generic method for diagnosis, discovery and sequencing of viruses. Virology. 2009;388: 1–7. doi:10.1016/J.VIROL.2009.03.024

18. Ma M, Huang Y, Gong Z, Zhuang L, Li C, Yang H, et al. Discovery of DNA Viruses in Wild-Caught Mosquitoes Using Small RNA High throughput Sequencing. PLoS One. 2011;6: e24758. doi:10.1371/JOURNAL.PONE.0024758

19. Tokarz R, Williams SH, Sameroff S, Sanchez Leon M, Jain K, Lipkin WI. Virome Analysis of Amblyomma americanum, Dermacentor variabilis, and Ixodes scapularis Ticks Reveals Novel Highly Divergent Vertebrate and Invertebrate Viruses. J Virol. 2014;88: 11480. doi:10.1128/JVI.01858-14

20. Aguiar E, Olmo RP, Marques JT. Virus-derived small RNAs: molecular footprints of host-pathogen interactions. WIREs RNA. 2016;7: 824–837. doi:10.1002/wrna.1361

21. Abbo SR, De Almeida JPP, Olmo RP, Balvers C, Griep JS, Linthout C, et al. The virome of the invasive Asian bush mosquito Aedes japonicus in Europe. Virus Evol. 2023;9. doi:10.1093/VE/VEAD041

22. Olmo RP, Todjro YMH, Aguiar ERGR, de Almeida JPP, Ferreira F V., Armache JN, et al. Mosquito vector competence for dengue is modulated by insect-specific viruses. Nat Microbiol. 2023;8: 135–149. doi:10.1038/s41564-022-01289-4

23. Kingston ER, Bartel DP. Ago2 protects Drosophila siRNAs and microRNAs rom target-directed degradation, even in the absence of 2-O-methylation. RNA. 2021;27: 710–724. doi:10.1261/RNA.078746.121/-/DC1

24. Kurth HM, Mochizuki K. 2′-O-methylation stabilizes Piwi-associated small RNAs and ensures DNA elimination in Tetrahymena. RNA. 2009;15: 675–685. doi:10.1261/RNA.1455509

25. Xiong Q, Zhang Y. Small RNA modifications: regulatory molecules and potential applications. Journal of Hematology & Oncology 2023 16:1. 2023;16: 1–24. doi:10.1186/S13045-023-01466-W

26. Korbel JO, Stegle O. Effects of the COVID-19 pandemic on life scientists. Genome Biol. 2020;21: 1–5. doi:10.1186/S13059-020-02031-1

27. Main BJ, Marcantonio M, Spencer Johnston J, Rasgon JL, Titus Brown C, Barker CM. Whole-genome assembly of Culex tarsalis. G3 Genes|Genomes|Genetics. 2021;11. doi:10.1093/G3JOURNAL/JKAA063

28. Baril C, Cassone BJ. Metatranscriptomic analysis of common mosquito vector species in the Canadian Prairies. mSphere. 2024;9. doi:10.1128/MSPHERE.00203-24

29. Faizah AN, Kobayashi D, Isawa H, Amoa-Bosompem M, Murota K, Higa Y, et al. Deciphering the virome of culex vishnui subgroup mosquitoes, the major vectors of japanese encephalitis, in Japan. Viruses. 2020;12. doi:10.3390/v12030264

30. Palatini U, Alfano N, Carballar RL, Chen XG, Delatte H, Bonizzoni M. Virome and nrEVEome diversity of Aedes albopictus mosquitoes from La Reunion Island and China. Virol J. 2022;19: 1–12. doi:10.1186/S12985-022-01918-8/FIGURES/5

31. Abel SM, Hong Z, Williams D, Ireri S, Brown MQ, Su T, et al. Small RNA sequencing of field Culex mosquitoes identifies patterns of viral infection and the mosquito immune response. Scientific Reports 2023 13:1. 2023;13: 1–13. doi:10.1038/s41598-023-37571-6

32. Batson J, Dudas G, Haas-Stapleton E, Kistler AL, Li LM, Logan P, et al. Single mosquito metatranscriptomics identifies vectors, emerging pathogens and reservoirs in one assay. Elife. 2021;10. doi:10.7554/ELIFE.68353

33. Rückert C, Prasad AN, Garcia-Luna SM, Robison A, Grubaugh ND, Weger-Lucarelli J, et al. Small RNA responses of Culex mosquitoes and cell lines during acute and persistent virus infection. Insect Biochem Mol Biol. 2019;109: 13–23. doi:10.1016/J.IBMB.2019.04.008

34. Siu RWC, Fragkoudis R, Simmonds P, Donald CL, Chase-Topping ME, Barry G, et al. Antiviral RNA Interference Responses Induced by Semliki Forest Virus Infection of Mosquito Cells: Characterization, Origin, and Frequency-Dependent Functions of Virus-Derived Small Interfering RNAs. J Virol. 2011;85: 2907–2917. doi:10.1128/JVI.02052-10

35. Czech B, Hannon GJ. One Loop to Rule Them All: The Ping-Pong Cycle and piRNA-Guided Silencing. Trends Biochem Sci. 2016;41: 324–337. doi:10.1016/J.TIBS.2015.12.008

36. Santos D, Feng M, Kolliopoulou A, Taning CNT, Sun J, Swevers L. What Are the Functional Roles of Piwi Proteins and piRNAs in Insects? 2023 [cited 18 Mar 2025]. doi:10.3390/insects14020187

37. Miesen P, Girardi E, Van Rij RP. Distinct sets of PIWI proteins produce arbovirus and transposon-derived piRNAs in Aedes aegypti mosquito cells. Nucleic Acids Res. 2015;43: 6545–6556. doi:10.1093/NAR/GKV590

38. Ozata DM, Gainetdinov I, Zoch A, O’Carroll D, Zamore PD. PIWI-interacting RNAs: small RNAs with big functions. Nature Reviews Genetics 2018 20:2. 2018;20: 89–108. doi:10.1038/s41576-018-0073-3

39. Altinli M, Leggewie M, Schulze J, Gyanwali R, Badusche M, Sreenu VB, et al. Antiviral RNAi Response in Culex quinquefasciatus-Derived HSU Cells. Viruses. 2023;15: 436. doi:10.3390/V15020436/S1

