## Supplementary material for "Virome profiling of *Culex tarsalis* through small RNA-seq: A challenge of suboptimal samples"

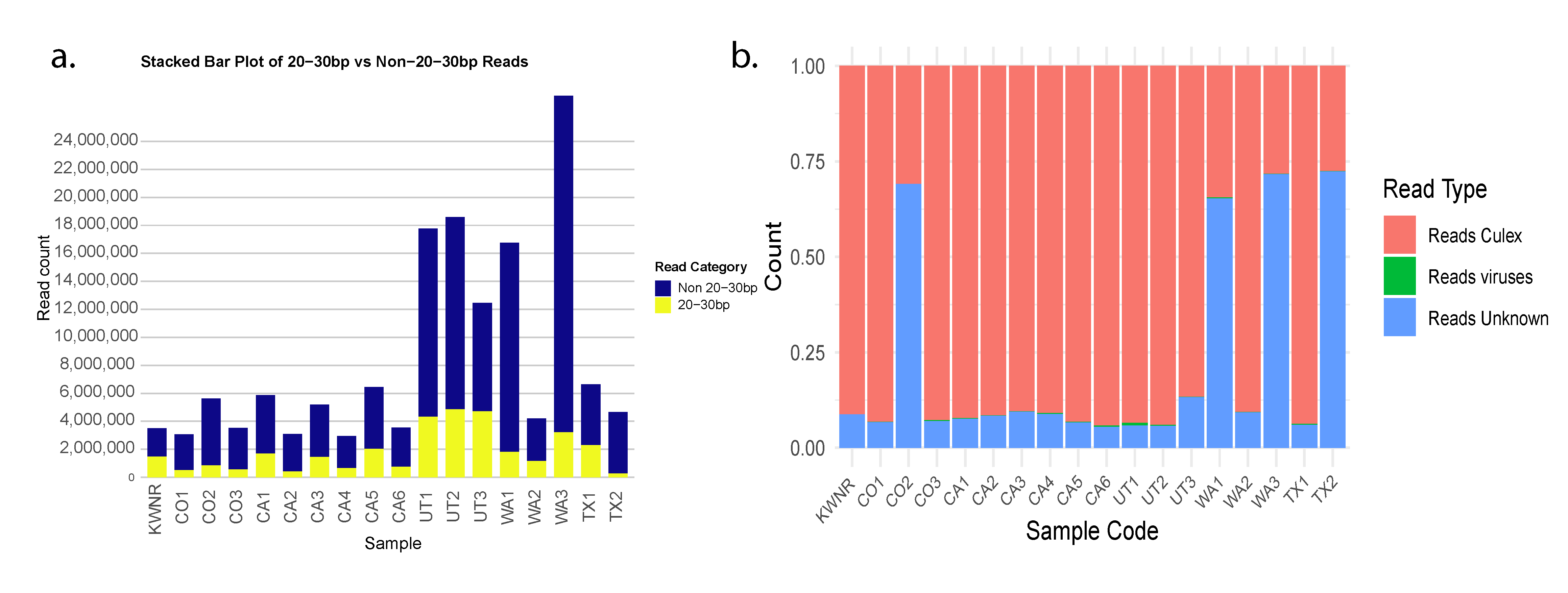

Supplementary figure 1. Library size and reads classification per location. Only reads that passed QC (processed reads) were used for plots.

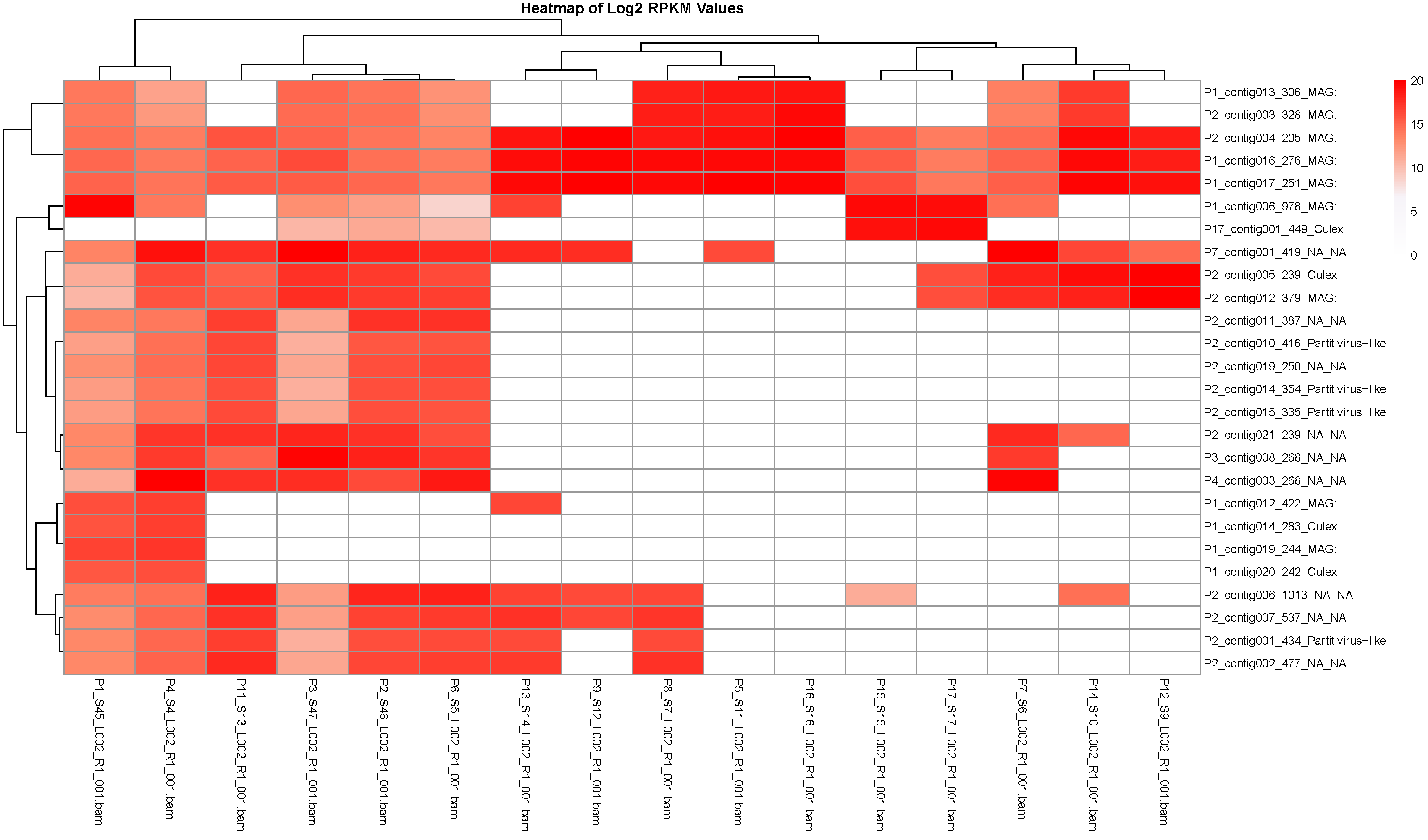

Supplementary figure 2. Heatmap and clustering of representative contigs. Libraries were mapped against the representative contigs and the number of reads mapped per contig was used to calculate the Log2 RPKM values. Library clusters, and their corresponding contigs, were used for contig extension.
